# cf-Cabernet: Nick Repair Prior to End Repair Preserves Endogenous DNA Methylation Information in Cell-Free DNA Methylation Sequencing

**DOI:** 10.64898/2026.07.14.738208

**Authors:** Tianjiao Yuan, Yali Bai, Liyang Song, Aicui Zhang, Yueyang Liu, Yunlong Cao

## Abstract

Cell-free DNA (cfDNA) methylation profiling holds great promise for non-invasive cancer detection, yet accurate methylome analysis is compromised by DNA damage inherent to cfDNA. Standard library preparation workflows involve an end-repair step during which DNA polymerases can initiate synthesis from single-strand breaks (nicks), replacing endogenous methylated nucleotides with unmethylated nucleotides in the 3′ direction and systematically erasing methylation information. This artifact is distinct from the terminal jagged-end effect and disproportionately affects cfDNA and FFPE DNA, which harbor abundant nicks. Here, we developed cf-Cabernet, which builds upon the Cabernet framework (Cao et al., 2023) — an enzymatic methylation sequencing method featuring carrier DNA-assisted sample recovery and amplification-friendly post-conversion processing — with the addition of a Taq DNA ligase-mediated nick repair step prior to end repair. Using matched cfDNA samples, we compared cf-Cabernet against standard EM-seq and WGBS. cf-Cabernet and EM-seq both substantially outperformed WGBS in alignment rate. Critically, while standard EM-seq exhibited globally reduced methylation levels compared to WGBS, cf-Cabernet yielded methylation levels concordant with WGBS. M-bias analysis revealed that EM-seq libraries showed persistently depressed methylation across the entire read length, whereas cf-Cabernet methylation recovered to the WGBS baseline beyond the terminal ∼40 bp jagged-end region. Nick-induced methylation erasure during end repair is a significant but previously underappreciated source of error in cfDNA methylation sequencing. cf-Cabernet effectively mitigates this artifact through pre-emptive nick ligation, enabling accurate methylome profiling from damaged DNA templates. This method is broadly applicable to cfDNA, FFPE DNA, and other clinical specimens where DNA integrity is compromised, providing a robust foundation for methylation-based liquid biopsy applications.

## Introduction

DNA methylation at cytosine-phosphate-guanine (CpG) dinucleotides is one of the most extensively studied epigenetic modifications, playing fundamental roles in transcriptional regulation, genomic imprinting, X-chromosome inactivation, and the maintenance of genomic stability[1,2]. Aberrant DNA methylation patterns, particularly hypermethylation of tumor suppressor gene promoters and global hypomethylation of repetitive elements, are hallmark features of human cancers and occur early during malignant transformation[3,4]. Consequently, analysis of cell-free DNA (cfDNA) methylation in peripheral blood has emerged as a promising strategy for non-invasive cancer detection, monitoring, and prognosis assessment[5,6]. The ability to accurately profile the methylome from cfDNA holds tremendous potential for liquid biopsy applications, yet the technical challenges inherent to cfDNA analysis remain substantial barriers to clinical translation.

For decades, bisulfite conversion-based methods, most notably whole-genome bisulfite sequencing (WGBS), have served as the gold standard for DNA methylation analysis at single-base resolution[7]. Despite its widespread use, bisulfite treatment suffers from several well-documented limitations. The harsh chemical conditions required for bisulfite conversion — high temperature combined with low pH and high bisulfite concentration — cause extensive DNA degradation through depurination and strand breakage[8,9]. This degradation is particularly problematic for cfDNA, which is already highly fragmented (∼166 bp modal size) and typically available in limited quantities[10]. The resulting loss of template molecules reduces library complexity, compromises alignment rates, and can introduce significant coverage bias. Moreover, bisulfite conversion cannot distinguish 5-methylcytosine (5mC) from 5-hydroxymethylcytosine (5hmC), an oxidized derivative with distinct biological functions[11].

To overcome these limitations, the enzymatic methyl-seq (EM-seq) method was developed as a bisulfite-free alternative that employs enzymatic conversion of unmethylated cytosines[12]. EM-seq uses TET2 and T4-β-glucosyltransferase (T4-BGT) to oxidize and protect 5mC and 5hmC, followed by APOBEC3A-mediated deamination of unprotected cytosines to uracils. This enzymatic approach preserves DNA integrity, resulting in higher library yields, improved alignment rates, and more uniform genomic coverage compared to WGBS[12,13].

Building upon the EM-seq framework, the Cabernet method (Carrier-Assisted Base-conversion by Enzymatic ReactioN with End-Tagging) was recently developed for single-cell methylation sequencing and demonstrated several key innovations that markedly improve DNA recovery and workflow efficiency[14]. Cabernet introduces carrier double-stranded DNA during the purification steps to prevent irreversible surface adsorption of picogram-level template DNA — a critical advance for low-input applications. Furthermore, Cabernet streamlines the post-conversion workflow by performing PCR amplification directly after APOBEC3A deamination without an intervening purification step, eliminating an additional source of DNA loss. These features collectively enable highly sensitive methylome profiling from minute quantities of DNA, achieving single-cell resolution in the original application[14]. The Cabernet framework — carrier-assisted recovery and direct post-conversion PCR — is directly applicable to cfDNA methylation analysis, where input amounts are often below 10 ng and DNA loss at each purification step can substantially compromise library complexity.

However, despite these advantages of enzymatic conversion-based methods, EM-seq, Cabernet, and other ligation-based library preparation methods share a fundamental, yet largely overlooked, limitation that disproportionately affects damaged DNA templates. During the end-repair step of standard library preparation, DNA polymerases with strand-displacement or nick-translation activity can initiate DNA synthesis from single-strand breaks (nicks) present in the template, replacing the original nucleotides downstream of the nick site in the 3′ direction with newly synthesized, unmethylated nucleotides[15]. This "nick-induced replacement" phenomenon effectively erases the endogenous methylation information from the affected strand segment. Critically, this issue is distinct from the well-characterized "jagged end" problem associated with the filling-in of 3′ recessed ends at the termini of double-stranded DNA fragments, which affects only the very ends of each molecule[16,17].

cfDNA and formalin-fixed paraffin-embedded (FFPE) tissue-derived DNA are particularly susceptible to this artifact because they contain abundant single-strand nicks. In cfDNA, nicks arise from multiple sources: endonucleolytic cleavage by caspase-activated DNase (CAD/DFFB) during apoptosis, further processing by circulating nucleases such as DNASE1L3 and DNASE1, oxidative DNA damage in the circulation, and mechanical shearing forces[10,18,19]. FFPE DNA accumulates nicks through formalin-induced crosslinking that strains the DNA helix, promoting apurinic sites and strand breaks, which are exacerbated by high-temperature decrosslinking during sample processing[20,21]. For methylation sequencing, the consequence is severe: every nick site in a template molecule serves as a potential initiation point for DNA polymerase during end repair, leading to the replacement of the original, potentially methylated, cytosines with unmodified cytosines along the entire 3′ direction from the nick. This can result in systematic underestimation of methylation levels across substantial portions of each affected read, with the magnitude of the effect proportional to the nick density and the distance from the nick site to the 3′ end of the fragment.

The extent of this artifact depends critically on the choice of DNA polymerase used during end repair. T4 DNA polymerase, which lacks both 5′→3′ exonuclease and strand-displacement activity, has been shown to preserve approximately 50% of original CpG methylation, whereas the Klenow fragment of DNA polymerase I results in 75–90% loss of methyl-cytosine signals[15]. Nevertheless, even with T4 polymerase, the residual nick-translation activity leads to measurable methylation erosion. For cfDNA, where nicks are abundant and input material is limited, any degree of methylation signal loss represents a significant analytical challenge — particularly when attempting to detect tumor-derived hypomethylation or hypermethylation at low variant allele frequencies.

Here, we present cf-Cabernet, which extends the Cabernet framework[14] to address the nick-induced methylation erasure problem in cfDNA. cf-Cabernet inherits the core innovations of Cabernet — carrier DNA-assisted sample recovery and post-APOBEC3A deamination direct PCR amplification — while adding a critical pre-end-repair step: treatment with Taq DNA ligase, an NAD⁺-dependent enzyme that specifically seals single-strand nicks in double-stranded DNA with high fidelity. This pre-treatment seals nicks before the DNA polymerase encounters them during end repair, effectively preventing polymerase-mediated replacement of methylated nucleotides. We systematically evaluated cf-Cabernet against standard EM-seq and WGBS using cfDNA samples, demonstrating that nick repair prior to end repair substantially preserves endogenous methylation signals, improves methylation calling accuracy, and eliminates the requirement for extensive read trimming beyond the terminal jagged-end region. Our approach provides a robust and accurate methylation profiling solution for cfDNA and other damaged DNA specimens, enabling more reliable liquid biopsy-based methylation analysis.

## Results

### Development of the cf-Cabernet workflow

The cf-Cabernet workflow builds upon the Cabernet framework[14] and introduces a critical nick-repair step prior to the enzymatic methylation sequencing library preparation pipeline (Figure 1). The procedure begins with the treatment of cfDNA using Taq DNA ligase, an NAD⁺-dependent ligase that specifically seals single-strand nicks in double-stranded DNA without ligating across gaps or joining blunt-ended fragments. This step repairs naturally occurring nicks in the cfDNA template, effectively removing the internal initiation sites that would otherwise be exploited by DNA polymerases during subsequent end repair. Following nick repair, the DNA undergoes standard end repair to generate blunt-ended fragments, after which methylated sequencing adapters are ligated to both ends. The adapter-ligated DNA is then subjected to enzymatic methyl-cytosine conversion using the EM-seq conversion module (TET2/T4-BGT oxidation and protection, followed by APOBEC3A deamination). Consistent with the Cabernet protocol[14], carrier double-stranded DNA is added prior to purification steps to minimize surface adsorption losses of the picogram-to-nanogram input cfDNA, and PCR amplification is performed directly after APOBEC3A deamination without an intervening column- or bead-based purification, thereby eliminating an additional source of template loss. The amplified library is then purified and subjected to high-throughput sequencing.

**Figure 1.**
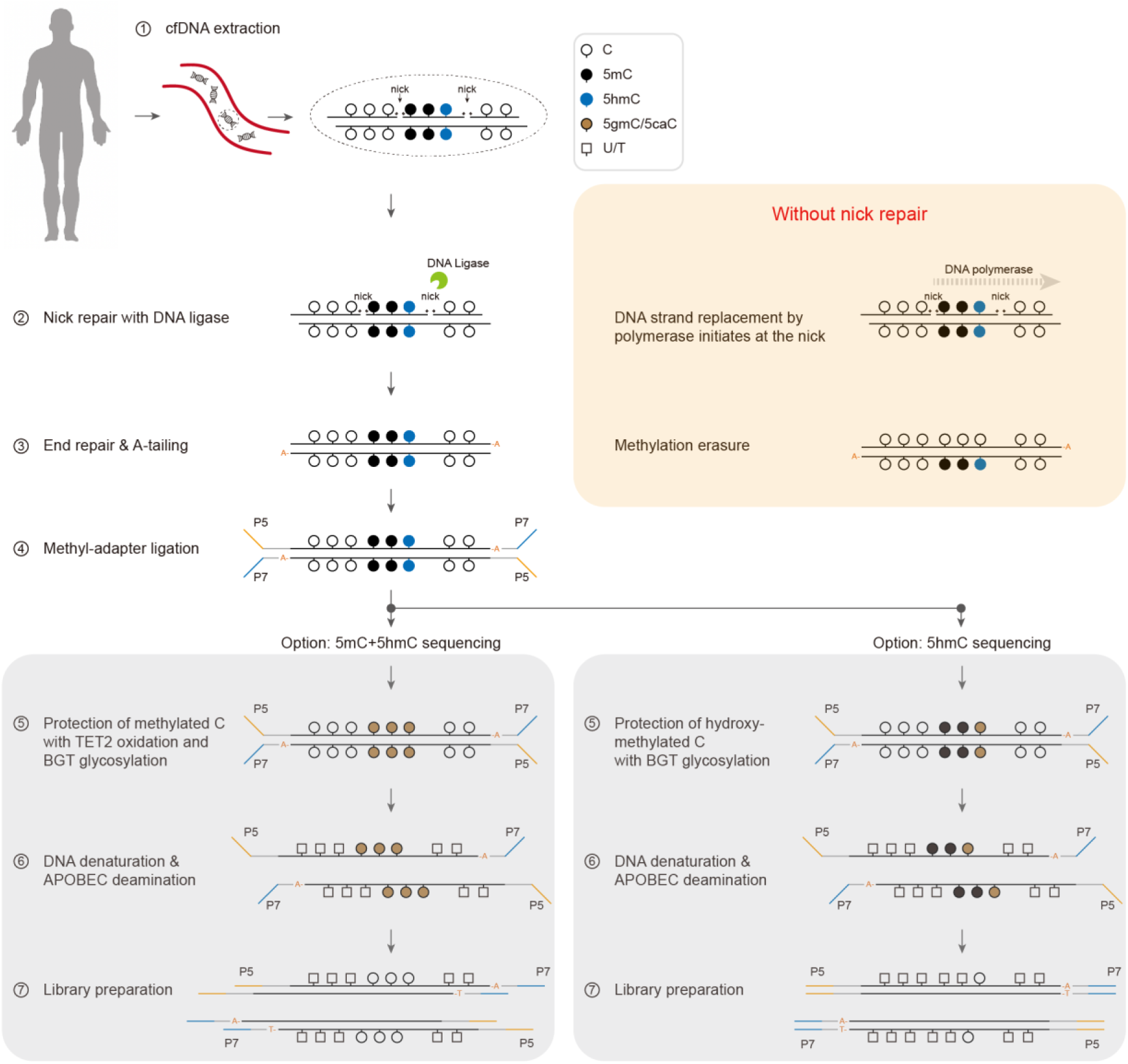
Schematic overview of the cf-Cabernet workflow. cfDNA is first treated with Taq DNA ligase to seal single-strand nicks, then undergoes end repair, methylated adapter ligation, and enzymatic conversion. Carrier DNA is added to prevent sample loss during purification, and PCR amplification is performed directly after APOBEC3A deamination without an intervening purification, following the Cabernet protocol.

### cf-Cabernet improves alignment rate and preserves global methylation levels

To evaluate the performance of cf-Cabernet, we sequenced cfDNA from the same sample using all three methods: WGBS, standard EM-seq, and cf-Cabernet. We first assessed the efficiency of C-to-T conversion in each method using spike-in control DNAs: unmethylated lambda DNA as a negative conversion control and fully methylated pUC19 DNA as a positive conversion control. cf-Cabernet achieved high conversion efficiencies on both control templates, with >99% of unmethylated cytosines converted in the lambda negative control and >99% of methylated cytosines protected in the pUC19 positive control (Figure 2A), confirming that neither the nick repair step nor the Cabernet-specific workflow modifications compromised conversion performance. We next compared alignment rates across methods. Both EM-seq and cf-Cabernet demonstrated substantially higher alignment rates compared to WGBS, consistent with the known advantage of enzymatic conversion in preserving DNA integrity and library complexity (Figure 2B). This improvement is particularly meaningful for cfDNA applications, where input material is inherently limited and maximizing usable sequencing reads is critical for detection sensitivity. The superior preservation of DNA integrity by enzymatic methods was further corroborated by fragment size distribution analysis: WGBS libraries showed a marked depletion of longer insert fragments relative to the input cfDNA, consistent with bisulfite-induced degradation preferentially breaking longer molecules, whereas EM-seq and cf-Cabernet libraries faithfully retained the full fragment size range of the input material (Figure 2C).

**Figure 2.**
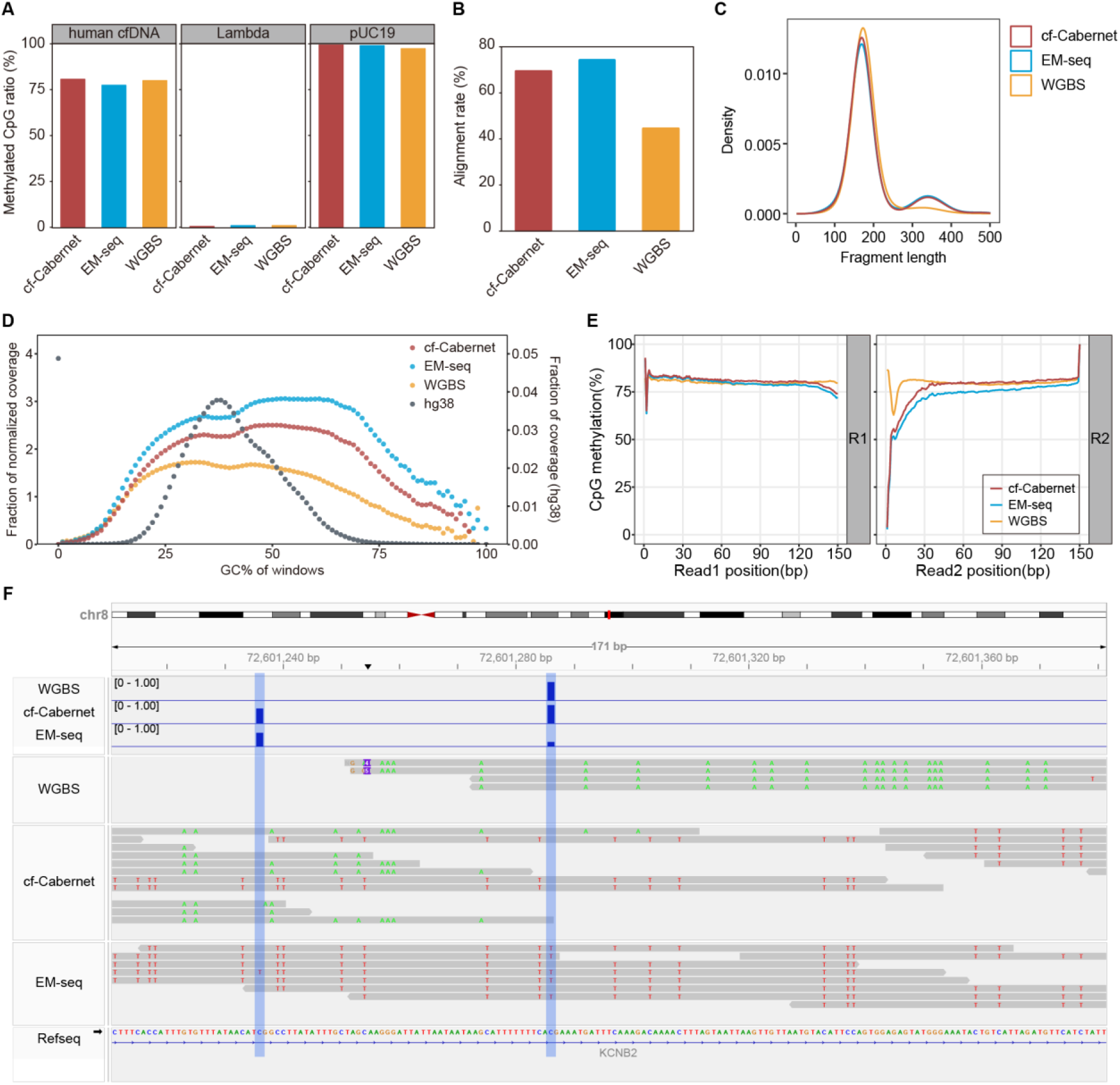
Library quality and methylation performance comparison of WGBS, EM-seq, and cf-Cabernet. **(A)** Alignment rates for each method. **(B)** Global CpG methylation ratios. **(C)** fragment size distribution of libraries showing that WGBS selectively loses longer fragments due to bisulfite-induced degradation, whereas EM-seq and cf-Cabernet faithfully retain the input fragment size range. **(D)** Normalized read coverage across genomic bins of varying GC content. WGBS shows pronounced depletion of high-GC regions and enrichment of low-GC regions, whereas EM-seq and cf-Cabernet exhibit substantially more uniform coverage. **(E)** M-bias plots showing methylation level as a function of position along Read 1 and Read 2 (nt 1–150). The cf-Cabernet library shows methylation recovery to the WGBS baseline beyond position 40, while EM-seq exhibits persistently depressed methylation across the entire read. **(F)** IGV snapshot of read-level methylation tracks at a representative genomic locus. WGBS and cf-Cabernet tracks show consistent methylation across individual reads at the highlighted CpG sites, whereas the EM-seq track displays a marked loss of methylation signal, with most individual reads reporting an unmethylated status.

We further examined GC bias — a well-documented limitation of WGBS arising from the DNA-degrading nature of bisulfite treatment. As previously characterized by Olova et al., the harsh bisulfite conditions induce an extra round of DNA fragmentation, preferentially destroying fragments derived from unmethylated C-rich genomic regions[9]. These degraded C-rich fragments are excluded from the library pool before PCR amplification, resulting in their systematic underrepresentation in the final sequencing data. To assess coverage uniformity, we partitioned the genome into bins by GC content and compared the normalized read coverage across methods. As expected, WGBS exhibited a pronounced coverage bias, with high-GC regions, such as CpG islands, showing substantially reduced coverage relative to intermediate-GC regions (Figure 2D). In contrast, both EM-seq and cf-Cabernet generated markedly more uniform coverage distributions across the full GC content spectrum. Although enzymatic conversion also deaminates unmethylated cytosines to uracils, following the same C-to-U conversion logic as bisulfite, the enzymatic reaction conditions are far milder and do not cause the extensive DNA strand breakage that selectively eliminates C-rich fragments from bisulfite-treated libraries.

We next compared the global CpG methylation levels measured by each method. WGBS and cf-Cabernet yielded closely concordant global methylation ratios, consistent with their shared ability to capture the endogenous methylation landscape. In striking contrast, standard EM-seq produced noticeably lower global methylation levels (Figure 2B). This discrepancy suggests that standard EM-seq, despite its advantages in DNA preservation, systematically underestimates methylation, particularly in templates harboring single-strand discontinuities.

To investigate the mechanistic basis of this observation, we performed M-bias analysis, which examines how methylation levels vary along the sequencing read (Figure 2E). In the cf-Cabernet library, the methylation levels on Read 2 showed a characteristic depression in the first ∼40 bp, consistent with the well-established jagged-end effect resulting from end-repair filling of 3′ recessed ends at fragment termini[16]. However, beyond position 40, cf-Cabernet methylation levels recovered fully and remained essentially identical to the WGBS baseline across the remainder of the read (positions 40–150). In sharp contrast, standard EM-seq libraries exhibited consistently reduced methylation levels across the entire length of Read 2, with no recovery to the WGBS baseline even at the innermost positions of the read. This persistent methylation deficit in EM-seq, extending far beyond the terminal jagged-end region, provides direct evidence that internal nick sites throughout the template molecule serve as additional DNA synthesis initiation points during end repair, leading to extensive replacement of methylated cytosines with unmethylated nucleotides along the full 3′ direction from each nick. The Taq ligase pre-treatment in cf-Cabernet effectively seals these internal nicks, confining end-repair-mediated replacement to the terminal jagged-end region only.

To directly visualize the impact of nick-induced methylation erasure at single-molecule resolution, we examined individual CpG sites using IGV, displaying aligned read-level methylation tracks from the same cfDNA sample processed by all three methods (Figure 2F). At CpG sites where both WGBS and cf-Cabernet consistently reported methylated cytosines across individual reads, the EM-seq reads exhibited a striking and widespread loss of methylation signal. This read-level methylation dropout occurred throughout the length of the sequencing reads, not merely at the terminal regions, consistent with the M-bias profile showing persistent undermethylation across the entire read. Notably, the cf-Cabernet track closely recapitulated the methylation pattern observed in WGBS at these same CpG positions, confirming that the nick repair step successfully preserved the endogenous methylation marks that were otherwise erased during standard EM-seq library preparation. This direct read-level comparison provides compelling visual evidence that single-strand nicks in cfDNA serve as widespread initiation points for polymerase-mediated strand replacement during end repair, effectively converting originally methylated CpG sites to unmethylated calls across the entire affected read.

### cf-Cabernet correctly capitulates methylation landscape across genomic features

To uniformly mitigate the confounding effect of jagged ends on methylation quantification, all subsequent EM-seq and cf-Cabernet analyses were performed after trimming the first 40 bp from the 5′ end of Read 2 for each read pair. Specifically, for each paired-end fragment, the 40 bp segment proximal to the Read 2 start site, corresponding to the 3′ terminus of the original cfDNA fragment, where end-repair-mediated filling-in of recessed ends introduces unmethylated nucleotides, was excluded from methylation calling. This trimming was applied to every fragment uniformly, ensuring that terminal jagged-end artifacts were systematically removed before any downstream methylation comparisons.

To quantitatively assess the concordance between methods, we computed pairwise Pearson correlations of CpG methylation levels across 100-kb genomic windows for all three datasets (Figure 3A). At this resolution, all three methods showed broadly similar methylation profiles, with pairwise correlation coefficients ranging from 0.83 to 0.87, indicating that the overall methylome architecture was faithfully captured by each technique. However, the correlation was slightly but consistently lower for the EM-seq versus WGBS comparison compared with the cf-Cabernet versus WGBS comparison (Figure 3A). This modest difference reflects the genome-wide accumulation of nick-induced methylation loss in EM-seq, which, while diluted at the 100-kb bin level, remains detectable as a systematic deviation from the WGBS reference. We note that the absolute correlation values were influenced by the relatively shallow sequencing depth typical of cfDNA methylome studies; deeper sequencing would be expected to further resolve the differences between methods by reducing sampling noise at individual CpG sites.

**Figure 3.**
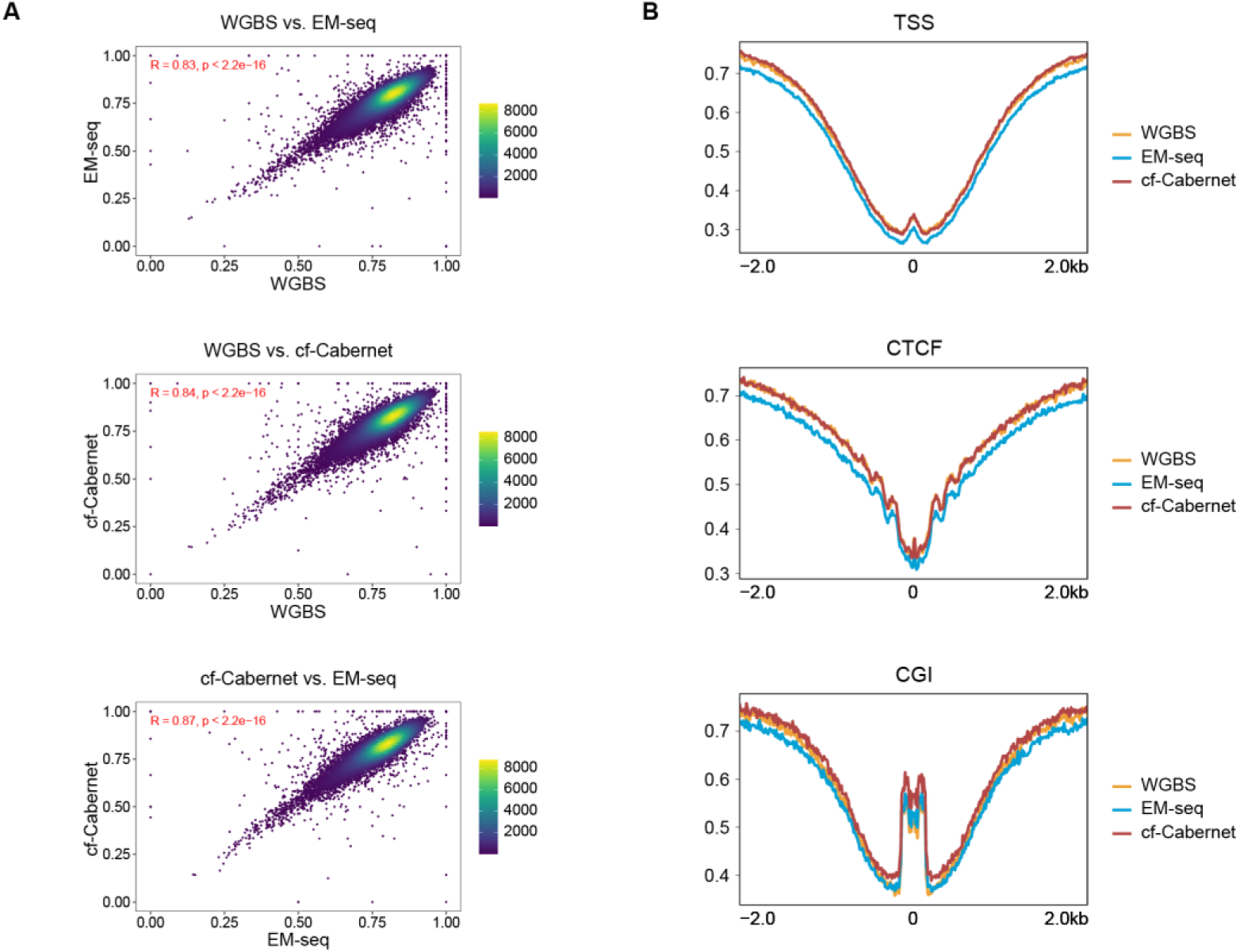
Methylation concordance and genomic feature-level validation of cf-Cabernet. **(A)** Pairwise Pearson correlation of CpG methylation levels in 100-kb genomic windows between WGBS, EM-seq, and cf-Cabernet. All three methods show broadly similar methylome profiles, with cf-Cabernet exhibiting higher correlation with WGBS than EM-seq does. **(B)** Mean CpG methylation profiles around transcription start sites (TSS), CTCF binding sites, and CpG islands (CGI). EM-seq shows systematically reduced methylation levels across all three feature categories, while cf-Cabernet closely tracks the WGBS profile.

To determine whether nick-induced methylation erasure preferentially affects specific genomic contexts, we profiled mean CpG methylation levels around transcription start sites (TSS), CTCF binding sites, and CpG islands (CGI) in all three datasets (Figure 3B). Consistent with the global methylation and M-bias observations, EM-seq exhibited noticeably lower methylation levels compared to both WGBS and cf-Cabernet across all three feature categories. In contrast, cf-Cabernet methylation profiles closely tracked the WGBS profiles at TSS, CTCF sites, and CGI regions, with near-identical mean methylation values and similar patterns of methylation enrichment and depletion around each feature (Figure 3B). These results demonstrate that the nick repair step in cf-Cabernet effectively protects methylation information across the regulatory genomic features relevant to epigenetic analysis, and that the protective effect is particularly critical for CpG-dense regions where the cumulative impact of nick-induced replacement is greatest.

### cf-Cabernet preserves native cfDNA fragment size distribution

Fragment size distribution is a foundational quality metric for cfDNA sequencing libraries. Bisulfite-based methods are known to preferentially degrade longer DNA molecules, but the extent to which they also affect shorter fragments, the predominant size class in cfDNA, has been less thoroughly characterized. To systematically evaluate fragment size representation, we compared the insert size distributions of cf-Cabernet, WGBS, and conventional whole-genome sequencing (WGS) libraries prepared from CRC patient-derived and healthy donor-derived cfDNA samples — using WGS as an orthogonal reference that captures the true fragment size profile without any C-to-T conversion step.

We stratified fragments into five size bins (50–100 bp, 100–150 bp, 150–200 bp, 200–250 bp, and 250–300 bp) and calculated the proportion of total fragments falling into each bin for both CRC patient sample (Figure 4A) and healthy control sample (Figure 4B). cf-Cabernet and WGS produced concordant fragment length distributions across all size bins in both the CRC and healthy control samples. In stark contrast, WGBS libraries exhibited a marked and selective depletion of fragments shorter than 150 bp. In the CRC patient sample, the 50–100 bp and 100–150 bp bins were underrepresented in WGBS (Figure 4A). A similar pattern was observed in the healthy control, with WGBS consistently showing reduced representation of the sub-150 bp fragment population (Figure 4B). This finding reveals that bisulfite treatment disproportionately eliminates short cfDNA fragments. The loss is likely attributable to the combined effects of bisulfite-mediated depurination and the subsequent alkaline treatment, which together can completely degrade already-fragmented short molecules, whereas longer molecules may sustain partial damage while retaining amplifiable segments.

**Figure 4.**
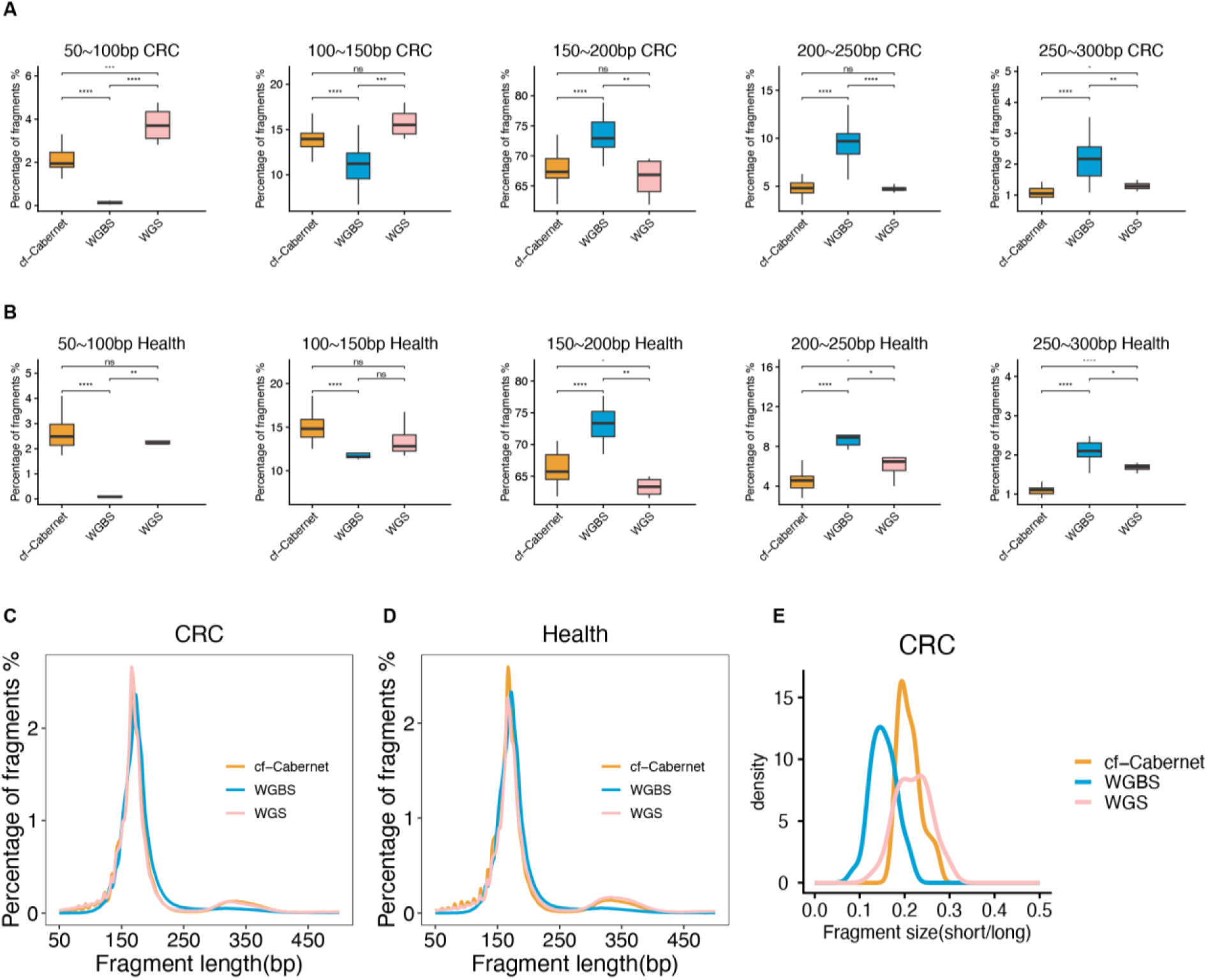
Fragment size preservation and fragmentomic fidelity of cf-Cabernet compared to WGBS and WGS. **(A,B)** Proportion of total fragments in each size bin (50–100 bp, 100–150 bp, 150–200 bp, 200–250 bp, 250–300 bp) for CRC patient sample (A) and healthy control sample (B). cf-Cabernet and WGS show highly concordant distributions, while WGBS selectively depletes fragments shorter than 150 bp. **(C, D)** Fragment size density distributions for CRC patient (C) and healthy control (D) cfDNA. cf-Cabernet and WGS curves are nearly superimposable, displaying the characteristic mononucleosomal (∼166 bp) and dinucleosomal (∼332 bp) peaks. WGBS exhibits a rightward shift with marked depletion of the short-fragment tail. **(E)** Density distribution of the short-to-long fragment ratio (<150 bp / >250 bp) in CRC patient cfDNA. cf-Cabernet and WGS produce closely overlapping distributions with nearly identical medians, whereas WGBS yields a substantially lower median short/long ratio.

We next visualized the fragment size density distributions for the CRC and healthy control samples (Figures 4C and 4D). WGS and cf-Cabernet both displayed the characteristic cfDNA fragment profile, with a dominant mononucleosomal peak at ∼166 bp and discernible dinucleosomal peaks at ∼332 bp. The density curves for these two methods were nearly superimposable across the entire size spectrum. WGBS, however, showed a pronounced rightward shift in the fragment size distribution: the short-fragment tail was substantially diminished relative to WGS and cf-Cabernet. The integrated density in the short-fragment region was visibly lower in WGBS, quantitatively consistent with the binned analysis in Figures 4A and 4B. Importantly, this distortion was observed in both the CRC and healthy control samples, indicating that it is a systematic method-intrinsic bias rather than a sample-specific artifact.

Given the clinical importance of fragmentomic features in liquid biopsy, we calculated the short-to-long fragment ratio for each library and compared the density distributions across methods (Figure 4E). The short/long ratio has emerged as a promising cfDNA-derived metric for cancer detection and monitoring, as tumor-derived ctDNA tends to be shorter than cfDNA from healthy cells, resulting in elevated short/long ratios in cancer patients relative to healthy individuals. cf-Cabernet and WGS yielded closely overlapping short/long ratio density distributions, with nearly identical median values. In contrast, WGBS produced a markedly shifted distribution, with a substantially lower median short/long ratio reflecting the selective loss of short fragments during bisulfite conversion (Figure 4E). This systematic depression of the short/long ratio in WGBS has direct clinical implications: by selectively depleting the short-fragment population that is preferentially enriched for tumor-derived DNA, WGBS may reduce the sensitivity of fragmentomics-based cancer detection. cf-Cabernet, by faithfully preserving the native cfDNA fragment size distribution, maintains the integrity of this clinically informative fragmentomic parameter.

## Discussion

Our study identifies and systematically characterizes a critical source of methylation detection bias in ligation-based methylation sequencing: the initiation of DNA polymerase-mediated strand replacement from internal single-strand nicks during the end-repair step. This phenomenon leads to progressive replacement of endogenous nucleotides — and consequently erasure of endogenous methylation marks — along the entire 3′ direction of each nick site. Unlike the well-recognized jagged-end effect that is confined to read termini, nick-induced replacement can affect the entire length of a sequencing read, representing a far more extensive source of methylation information loss.

We demonstrated that cfDNA, which is naturally enriched for single-strand breaks due to its apoptotic origin and exposure to nuclease activity in circulation, is especially vulnerable to this artifact. Standard EM-seq, despite its documented advantages over bisulfite-based methods in preserving DNA integrity, provides no protection against nick-induced methylation erasure and consequently produces systematically underestimated methylation levels across the full read length, as evidenced by our M-bias analysis. This finding has important implications for the interpretation of existing cfDNA methylation datasets generated using standard EM-seq or other ligation-based library preparation methods.

The cf-Cabernet method addresses this limitation through a conceptually simple yet mechanistically effective intervention: sealing nicks with Taq DNA ligase prior to the end-repair reaction. This pre-treatment confines polymerase-mediated replacement to the terminal jagged-end region, which can then be readily managed through standard read trimming. Across 100-kb genomic windows, cf-Cabernet achieves pairwise methylation correlation with WGBS that exceeds that of standard EM-seq, while maintaining the superior alignment rates and DNA preservation characteristic of enzymatic conversion. The protective effect of nick repair is particularly evident at regulatory genomic features — TSS, CTCF binding sites, and CpG islands — where cf-Cabernet methylation profiles closely mirror those of WGBS, in contrast to the systematically depressed levels observed in EM-seq.

Beyond methylation accuracy, our fragment size analysis revealed an additional and clinically significant advantage of enzymatic approaches over WGBS: the selective loss of short cfDNA fragments (<150 bp) during bisulfite conversion. These short fragments constitute a substantial fraction of total cfDNA and are of particular biological interest, as tumor-derived ctDNA is enriched in this size range. The observed depletion of short fragments in WGBS translates directly into a depressed short-to-long fragment ratio, a metric increasingly used for cancer detection and tissue-of-origin inference in liquid biopsy. By faithfully preserving the native fragment size distribution, cf-Cabernet maintains the integrity of fragmentomic features that are orthogonal and complementary to methylation-based biomarkers. This dual preservation — of both methylation information and fragment size distribution — positions cf-Cabernet as a uniquely suitable platform for integrative cfDNA analyses that combine epigenetic and fragmentomic readouts.

Several aspects of our approach merit further discussion. First, Taq DNA ligase was chosen for its strict specificity for nicked duplex DNA, as it requires perfect base-pairing at the ligation junction and an available 5′ phosphate group. It does not ligate across gaps or join blunt-ended fragments, thereby minimizing the risk of generating chimeric molecules[24]. Second, the nick repair step is performed prior to end repair without an intervening purification, maintaining a streamlined workflow compatible with high-throughput processing. Third, the 40 bp terminal trimming window we applied is consistent with the expected extent of the jagged-end effect in cfDNA, as previously characterized by Jiang et al.[16]. We note that this trimming parameter may require optimization for other sample types with different fragment end characteristics, such as FFPE DNA or urinary cfDNA.

Our work has several important implications for the field of cfDNA methylome. It provides a mechanistic explanation for the systematic methylation underestimation observed in some EM-seq datasets, particularly those derived from damaged or fragmented DNA templates. The cf-Cabernet workflow is straightforward to implement using commercially available reagents and can be integrated into existing methylation sequencing pipelines with minimal additional cost or hands-on time. Beyond cfDNA, the method is broadly applicable to any sample type harboring single-strand breaks — including FFPE DNA, ancient DNA, and forensic specimens — which would be expected to benefit from pre-end-repair nick ligation.

A limitation of our study is that, while we demonstrated the performance of cf-Cabernet in cfDNA from colorectal cancer, broader validation across additional cancer types and larger patient cohorts will be necessary to establish the generalizability of our findings. Future work should also explore the integration of cf-Cabernet with single-stranded library preparation methods[25], which may provide complementary benefits for capturing the most heavily damaged template molecules.

In conclusion, the cf-Cabernet workflow extends the Cabernet framework[14] to provide a practical and effective solution to the nick-induced methylation erasure problem in cfDNA methylation sequencing. Our method enables more accurate methylome profiling from damaged DNA templates. We anticipate that cf-Cabernet will support the development of robust methylation-based biomarkers for cancer detection, monitoring, and precision medicine.

## Methods

### cfDNA sample preparation

Blood samples were collected in EDTA or Streck tubes, and plasma was separated by centrifugation within 4 hours. cfDNA was extracted using the QIAamp Circulating Nucleic Acid Kit (Qiagen) according to the manufacturer’s instructions. cfDNA concentration and fragment size distribution were assessed using the Agilent Bioanalyzer High Sensitivity DNA Kit.

### cf-Cabernet library preparation

Extracted cfDNA was first treated with Taq DNA ligase (New England Biolabs) in 1× Taq DNA Ligase Reaction Buffer containing 1 mM NAD⁺ at 45°C for 15 minutes. Nick-repaired cfDNA was then subjected to end repair, dA-tailing, and methylated adapter ligation using the NEBNext Ultra II End Repair/dA-Tailing Module and NEBNext Ligation Module. Adapter-ligated DNA was enzymatically converted using the NEBNext Enzymatic Methyl-seq Conversion Module. Following the Cabernet protocol[14], carrier double-stranded DNA was added to the conversion reaction, and after APOBEC3A deamination, the reaction mixture was used directly as template for PCR amplification without an intervening purification step. Libraries were PCR-amplified for 8–12 cycles and purified with AMPure XP beads.

### Standard EM-seq and WGBS

Parallel aliquots were processed through standard EM-seq (without Taq ligase pre-treatment) following the NEBNext Enzymatic Methyl-seq Kit protocol. For WGBS, libraries were prepared using the Accel-NGS Methyl-Seq DNA Library Kit (Swift Biosciences) and bisulfite-converted using the EZ DNA Methylation-Gold Kit (Zymo Research).

### Sequencing and data processing

Libraries were sequenced on an Illumina NovaSeq 6000 platform (2 × 150 bp paired-end). Reads were trimmed using Trim Galore (--paired --clip_r1 10 --clip_r2 10 --three_prime_clip_r1 10 --three_prime_clip_r2 10 for WGBS; --clip_r2 40 --three_prime_clip_r2 40 for cf-Cabernet and EM-seq). Trimmed reads were aligned to the hg38 reference genome using Bismark (v0.23.0) with Bowtie 2. Methylation calls were extracted using bismark_methylation_extractor. M-bias plots and global methylation statistics were generated using MethylDackel. Pearson correlation analyses were performed on CpG sites with ≥5× coverage in all three datasets. IGV (v2.8) was used for visualization of methylation tracks at individual loci.

## Data Availability

Analysis scripts and processed methylation data are available from the corresponding author upon reasonable request.

## Acknowledgments

This work is financially supported by Changping Laboratory (2026C-01-01, Y.C.).

## Author contributions

Y.C. supervised the study. T.Y., Y.B., L.S., and Y.C. developed and optimized the cf-Cabernet. T.Y. generated the data. Y.B., T.Y., A.Z., and Y.L. preprocessed the data and performed bioinformatic analyses. T.Y., Y.B., and Y.C. wrote the manuscript. All authors edited and approved the manuscript.

